# A genome-wide CRISPR screen implicates plasma membrane asymmetry in exogenous C6-ceramide toxicity

**DOI:** 10.1101/2022.09.26.509629

**Authors:** Siti Nur Sarah Morris, Kirandeep K. Deol, Mike Lange, James A. Olzmann

## Abstract

The bioactive sphingolipid ceramide impacts diverse cellular processes (e.g., apoptosis and cell proliferation) through its effects on membrane dynamics and intracellular signalling pathways. The dysregulation of ceramide metabolism has been implicated in cancer evasion of apoptosis and targeting ceramide metabolism has potential therapeutic benefits as a strategy to kill cancer cells and slow tumor growth. However, the mechanisms of cancer cell resistance to ceramide-mediated cell death are vastly intertwined and incompletely understood. To shed light on this mystery, we performed a genome wide CRISPR-Cas9 screen to systematically identify regulators of cancer resistance to the soluble short chain ceramide, C6 ceramide (C6-Cer). Our results reveal a complex landscape of genetic modifiers of C6-Cer toxicity, including genes associated with ceramide and sphingolipid metabolism, vesicular trafficking, and membrane biology. Furthermore, we find that loss of the phospholipid flippase subunit TMEM30A impairs the plasma membrane trafficking of its binding partner the P4-type ATPase ATP11B, and depletion of TMEM30A or ATP11B disrupts plasma membrane asymmetry and promotes resistance to C6-Cer toxicity independent of alterations in C6-Cer uptake. Together, our findings provide a resource of genetic modifiers of C6-Cer toxicity and reveal an unexpected role of plasma membrane asymmetry in C6-Cer induced cell death.

## INTRODUCTION

Ceramides are a class of bioactive sphingolipids that consist of a sphingoid base amide bound to a fatty acid (For reviews on ceramide see (Chaurasia and Summers, 2021; Morad and Cabot, 2013; Summers et al., 2019)). The length and degree of saturation within the sphingoid base or the fatty acid determines the biological activities of the individual ceramides. Multiple pathways generate ceramide, including the breakdown of complex sphingolipids such as sphingomyelins by sphingomyelinases, the reacylation of sphingosine in the ceramide salvage pathway, and the *de novo* synthesis pathway involving the condensation of serine and palmitoyl-CoA.

Ceramide has been implicated in diverse cellular roles, including the regulation of apoptosis, autophagy, cell proliferation, immune response, and ER stress (Chaurasia and Summers, 2021; Morad and Cabot, 2013; Summers et al., 2019). These varied functions are mediated by ceramide’s ability to regulate membrane dynamics (e.g., membrane fluidity through ceramide-enriched membrane platforms) (Grassmé et al., 2007) and intracellular signal transduction (e.g., its interactions with a host of effector proteins) (Stancevic and Kolesnick, 2010; Summers et al., 2019). For example, in the extrinsic apoptotic pathway, initiation by activation of death receptors belonging to the tumor necrosis factor (TNF) receptor superfamily triggers an increase in plasma membrane ceramide levels through the actions of sphingomyelinases (Pettus et al., 2002). The increased ceramide forms ceramide-enriched membrane platforms in the plasma membrane that prime death receptor clustering, facilitating the formation of death-inducing signalling complexes (Grassmé et al., 2003; Stancevic and Kolesnick, 2010). Ceramide’s position in the apoptotic pathway makes it a desirable topic for cancer research as dysregulation of ceramide metabolism has been implicated in cancer resistance to apoptosis. For example, reduced ceramide is associated with resistance to CD95 and TRAIL-induced apoptosis in a variety of cancer cell types (Voelkel-Johnson et al., 2005; White-Gilbertson et al., 2009). Moreover, therapeutics targeting ceramide anabolic enzymes sensitize cancer cells to apoptosis inducers. Similarly, raising the levels of ceramide through exogenous treatment with soluble short chain ceramide such as C6 ceramide (C6-Cer) can induce apoptosis in cancer cells and in vivo experiments delivering C6-Cer through nanoliposomes have been efficacious in pre-clinical mouse models of cancer (Flowers et al., 2012; Ji et al., 2010; Liu et al., 2010; Stover et al., 2005; Tagaram et al., 2011; Tran et al., 2008). These findings suggest that targeting ceramide metabolism could be an effective strategy for cancer treatment.

The incomplete understanding of the mechanisms that mediate cancer resistance or sensitivity to ceramide-related cell death remains an obstacle for further development of targeted therapeutics. Here, we employ genome wide CRISPR-Cas9 screens to provide a resource of genetic modifiers that influence C6-Cer toxicity. Furthermore, our findings reveal that a phospholipid flippase composed of TMEM30A and the P4-type ATPase ATP11B is required for C6-Cer toxicity, implicating membrane asymmetry as a key factor in C6-Cer-induced cell death.

## RESULTS

### Genome-wide CRISPR-Cas9 screen identifies regulators of C6-Cer toxicity

K562 chronic myelogenous leukemia cells have been previously shown to be sensitive to C6-Cer induced cell death, providing a useful model to study mechanisms of ceramide toxicity (Morad et al., 2015; Nica et al., 2008). Consistent with previous findings, K562 cells were sensitive to C6-Cer induced cell death with an EC50 of 27.90 μM (**Figure 1A**). To systematically identify genetic modifiers of exogenous ceramide toxicity, we performed a genome wide CRISPR-Cas9 knockout (KO) screen (**Figure 1B**). K562 cells expressing Cas9 were infected with an sgRNA library containing 212,821 sgRNAs targeting 20,500 protein-coding genes (∼10 sgRNAs/gene) along with several thousand control sgRNAs. The cells were then treated with a vehicle or at a 50% lethal concentration of C6-Cer. Four rounds of C6-Cer treatments were performed (with time for recovery), allowing sgRNAs that confer protection to be enriched and sgRNAs that confer sensitization to be depleted relative to the vehicle treated controls. The frequencies of the sgRNAs in the untreated and C6-Cer treated cell populations were then determined by deep sequencing, and the enriched and depleted genes identified using Cas9 high-throughput maximum likelihood estimator (casTLE) (Morgens et al., 2016; Morgens et al., 2017). The screening procedure was performed in duplicate and the data from both sets of screens were combined into a volcano plot depicting the collective casTLE confidence scores versus the casTLE phenotype effect score (**Figure 1C**). Employing a 1% false discovery rate (FDR), we identified 97 genes that significantly influence C6-Cer toxicity, including genes that - when depleted - result in resistance (64 genes) and sensitization (33 genes) to C6-Cer toxicity (**Figure 1C** and **Supplementary Table S1**).

**Figure 1.**
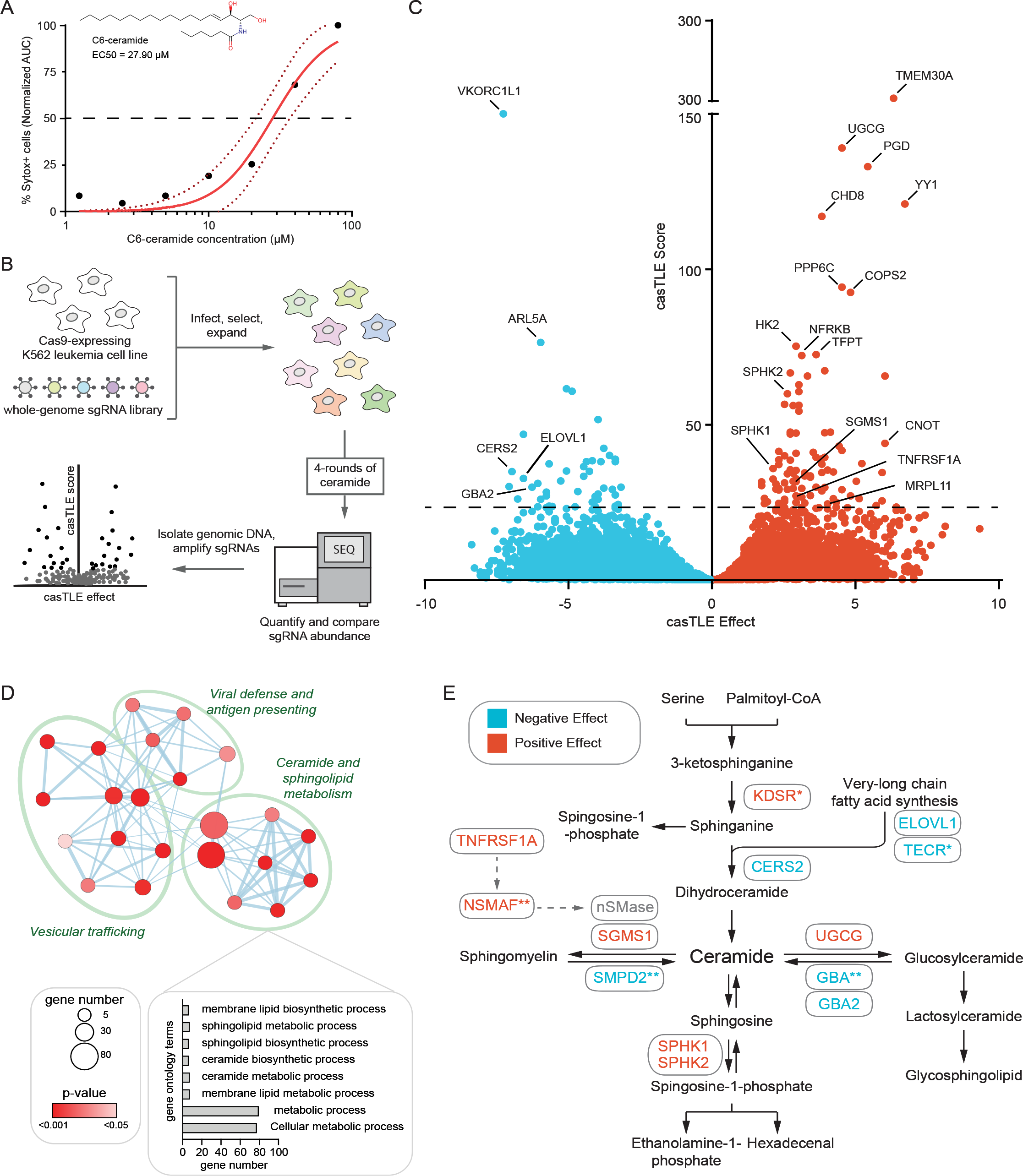
Genome-wide CRISPR-Cas9 screen identifies genetic modifiers of exogenous ceramide toxicity. A) K562 cells were treated with the indicated concentrations of C6-Cer and percentage Sytox Green positive (Sytox+) cells was measured using an IncuCyte over 24 hr. The structure of C6-Cer and the calculated EC50 are shown in the inset. Dotted lines indicate the 95% confidence intervals for the fitted curves. B) Schematic of the CRISPR-Cas9 screening strategy. C) Gene effect and gene score were calculated for individual genes analysed in the CRISPR-Cas9 screen using the CasTLE. D) Network map of enriched GO terms. Node size indicates gene number associated with a GO term and color indicates p-value. E) Ceramide metabolic network indicating genes identified in the screen and whether knockout has a negative (blue) or positive (red) effect on cell survival in response to sequential rounds of C6-Cer treatments.

Analysis of Gene Ontology (GO) pathway enrichments for the identified genetic modifiers revealed expected connections to ceramide and sphingolipid metabolism (**Figure 1D,E** and **Supplementary Figure 1A,B**). C6-Cer resistance factors (i.e., cells were sensitized to C6-Cer toxicity when these genes were depleted) that canonically involve ceramide metabolism included ceramide synthase, CERS2; the very-long chain fatty acid synthesis enzymes, ELOVL1 and TECR; sphingomyelin phosphodiesterase, SMPD2; and glucoceramidases, GBA and GBA2 (**Figure 1E**). C6-Cer sensitising factors (i.e., cells were resistant to C6-Cer death when these genes were depleted) included ceramide glucosyltransferase, UGCG; the sphingosine kinases, SPHK1 and SPHK2; and sphingomyelin synthase, SGMS1 (**Figure 1E**). Why the paradoxical depletion of genes associated with ceramide anabolism would result in resistance to C6-Cer toxicity is not immediately clear. It may be that these cell lines have adapted to higher levels of ceramides with increased protective mechanisms to suppress ceramide toxicity. In addition to genes directly involved in ceramide and sphingolipid metabolism, the TNF receptor TNFRSF1A (also known as TNFR1) and NSMAF were also identified as sensitizing factors (**Figure 1E**). The identification of ceramide and sphingolipid metabolic genes, as well as extrinsic apoptosis factors that act through ceramide, is consistent with the high performance of our CRISPR-Cas9 screening platform.

The GO analysis also revealed significant enrichments in genes involved in vesicular trafficking and membrane biology (**Figure 1D** and **Supplementary Figure 1A,B**). Consistent with this relationship, mapping genetic modifiers of C6-Cer toxicity onto a cell diagram identified a high number of genes within the secretory pathway. These included genes related to endoplasmic reticulum (ER) and Golgi trafficking as well as endocytic genes related to endosome and lysosome functions (**Figure 2**). The connection with vesicular trafficking is in part because several of the enzymes involved in ceramide and sphingolipid metabolism localize to the plasma membrane and thus, secretory pathways (**Figure 2**). However, many other genes involved in secretory protein trafficking were present, including Rab GTPases: Rab2A, Rab6A, Rab11a; the RAB GTPase activity proteins (RAB GAPs): TBC1D5 and ARF1; the cargo receptor, TMED2; the tSNARE protein, G0SR1; the EARP and GARP complex-interacting protein 1, TSSC1; the subunits of the clathrin-associated adaptor protein complex 1: AP1S1, AP1M1, AP1G1; and the clathrin-associated adaptor protein complex 2, AP2A1 (**Figure 2**). Many genes in the cell map were also present in nuclear transcriptional pathways (**Figure 2**), potentially reflecting the influence of multiple gene expression programs on ceramide and sphingolipid metabolism.

**Figure 2.**
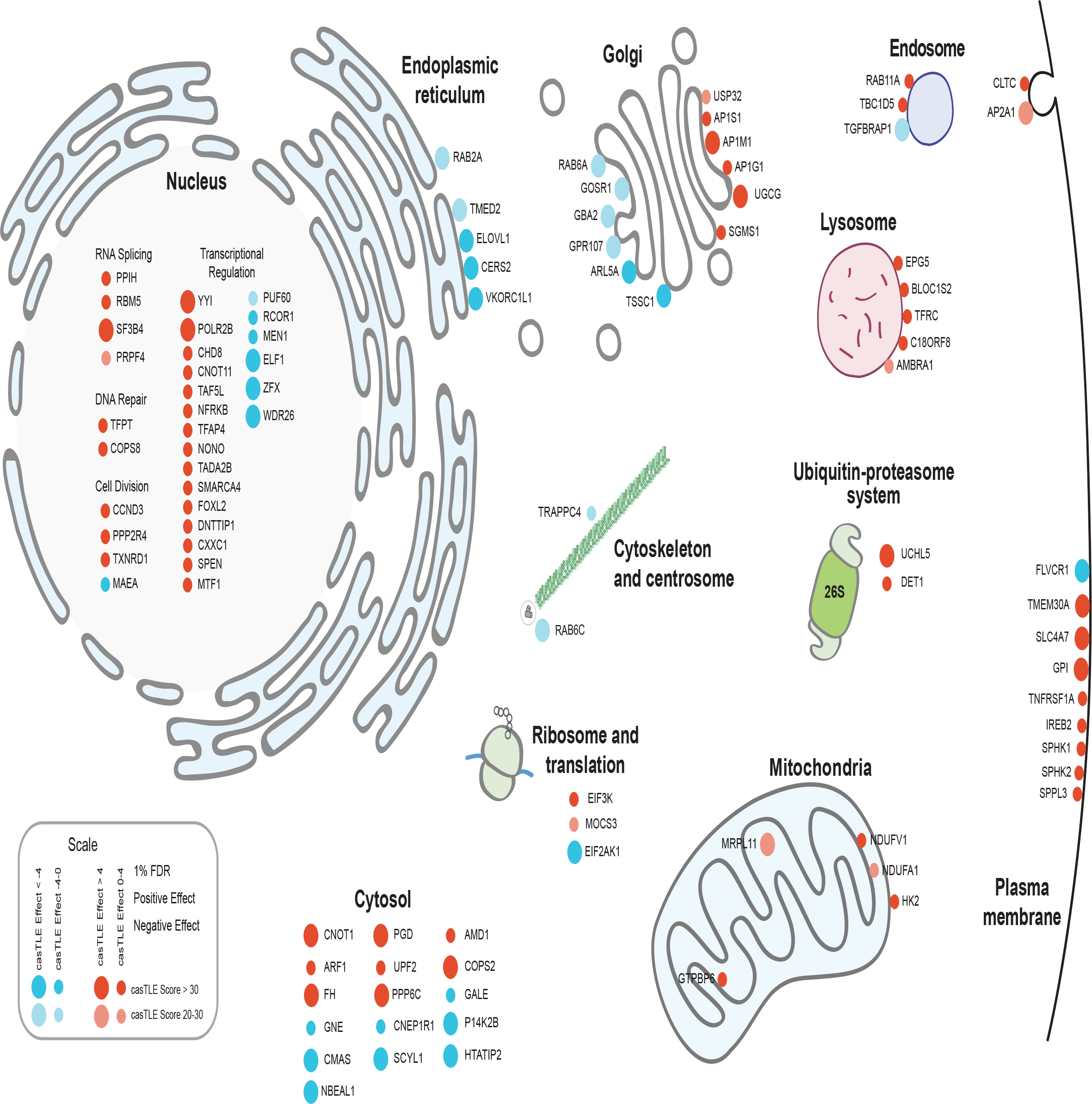
Cell map of genetic modifiers of C6-Cer toxicity identified in CRISPR-Cas9 screens. Cell diagram displaying XX of the most significant hits detected in genome-wide screens for regulators of C6-Cer toxicity. Node size indicates casTLE score (i.e., confidence) and color indicates casTLE effect (i.e., phenotype), with the positive effect in red and negative effect in cyan.

### Validation of select genetic modifiers of C6-Cer toxicity

We validated our screen results through competitive growth assays of 10 genes. Our selection criteria of the genes to validate were based upon their cellular functions (e.g., known involvement in cellular trafficking and membrane homeotasis) and the strength of the casTLE confidence and effect scores (within the 1% false-discovery rate). In this assay (**Figure 3A**), Cas9 expressing cell lines were infected with sgRNAs targeting candidate regulators of interest. Cells expressing the sgRNA plasmid co-express mCherry. These cells are combined in an equal ratio with a control line that does not express mCherry. This mixed cell population is treated with C6-Cer and then analyzed using flow cytometry to calculate the ratio of mCherry positive cells to mCherry negative cells. Cell lines expressing sgRNAs that confer resistance to C6-Cer death will be proportionally higher in number than the control cells, while cell lines expressing sgRNAs that sensitize them to C6-Cer will be less abundant than the control cells. Expression of sgRNAs targeting ceramide metabolic factors showed the anticipated effects based on our screen results, with depletion of CERS2 and GBA sensitizing cells to C6-Cer and depletion of UGCG promoting resistance to C6-Cer (**Figure 3B**). In addition, sgRNAs targeting CHD8 and TMEM30A both resulted in strong resistance to C6-Cer toxicity and sgRNAs targeting ARL5 sensitized cells to C6-Cer (**Figure 3B**). We did not observe a statistically significant effect of sgRNAs targeting MRPL11, VKORC1L1, CNOT1, or C18ORF8 (**Figure 3B**). This difference may be due to insufficient depletion of the target or due to the difference in treatment paradigms – in the screen, four consecutive C6-Cer treatments were performed versus the single treatment during the competitive growth assay.

**Figure 3.**
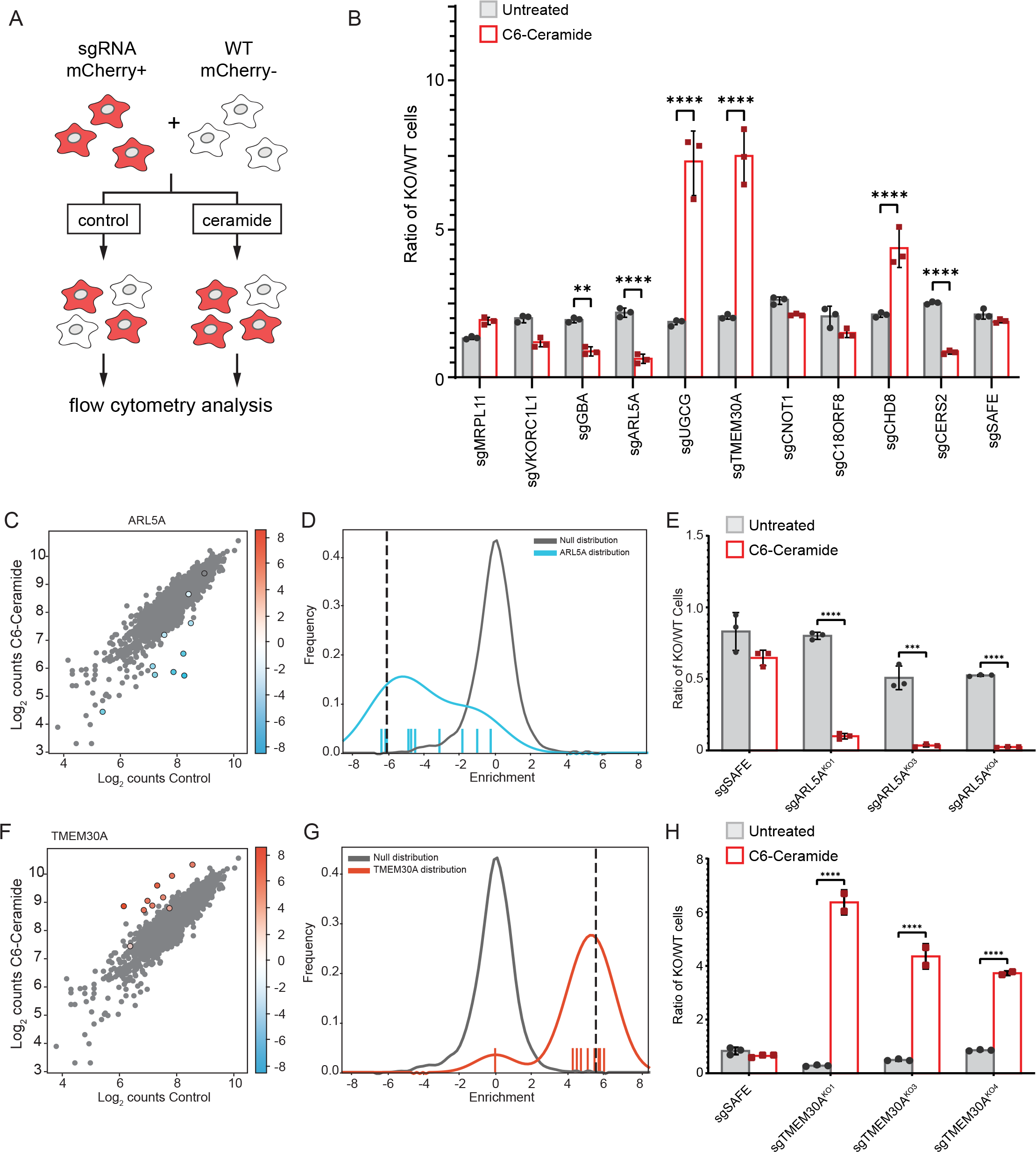
Validation of select regulators of exogenous C6-Cer toxicity. A) Schematic of fluorescent competitive growth assay. B) Ratio of KO/WT cells grown in the competitive growth assay depicted in panel A. Cells are untreated or treated with 30 μM C6-Cer for 24 hr and allowed to recover prior to analysis. C,D) Cloud plot and histogram indicating the disenriched sgRNAs targeting ARL5A following C6-Cer treatment in the CRISPR screen. E) Competitive growth assay for analysis of three independent ARL5A KO lines. F,G) Cloud plot and histogram indicating the enriched sgRNAs targeting TMEM30A following C6-Cer treatment in the CRISPR screen. H) Competitive growth assay for analysis of three independent TMEM30AKO lines.

Among the panel of genes analyzed, sgRNAs targeting ARL5A and TMEM30A yielded the strongest sensitization and resistance to C6-Cer, respectively (**Figure 3B**). KO of UDP-glucose ceramide glucosyltransferase (UGCG) also yielded strong resistance to C6-Cer. As UGCG is a gene that is known to be connected with ceramide metabolism, we chose to follow up on the more novel regulators – ARL6A and TMEM30A. Consistent with the high confidence identification of these candidate regulators, 8 of the 10 ARL5a-targeting sgRNAs were depleted and 10 of the 10 TMEM30A-targeting sgRNAs were enriched following C6-Cer treatment in our screen (**Figure 3C,D,F,G**). ARL5a is a small GTPase that may function in the recruitment of the GARP complex to the trans-Golgi network to regulate retrograde trafficking. TMEM30A is a subunit of phospholipid flippase complexes. It binds and facilitates the trafficking of several different P4-type ATPases to the plasma membrane, where they constitute functional lipid flippase complexes that regulate membrane asymmetry (Best et al., 2019; Hankins et al., 2015). To further validate these regulators of C6-Cer toxicity, we generated additional lines expressing unique sgRNAs targeting ARL5a and TMEM30A. Commercially available antibodies to our knockouts proved unreliable, therefore, we verified our lines through qPCR and Tracking of Indels by Decomposition (TIDE)^79^ (**Supplementary Figure 2**). Three independent ARL5a-targeting sgRNAs sensitized cells to C6-Cer (**Figure 3E**) and TMEM30A-targeting sgRNAs promoted resistance (**Figure 3H**) to C6-Cer in the competitive growth assay. Furthermore, we found that two TMEM30A knockout cell lines exhibited enhanced resistance to C6-Cer induced cell death in a flow cytometry assay employing SYTOX Green, a membrane impermeable cell death marker (**Supplementary Figure 3**). Together, these data establish ARL5a and TMEM30A as novel regulators of C6-Cer toxicity.

### TMEM30A disrupts membrane asymmetry, but does not influence C6-Cer flipping or uptake

Phospholipid flippase complexes are composed of a catalytic α-subunit – which can be one of several P4-type ATPases – and the single accessory β-subunit TMEM30A (Best et al., 2019; Coleman and Molday, 2011; Shin and Takatsu, 2019; Timcenko et al., 2019). The association of TMEM30A with the P4-type ATPase enables the ER exit and trafficking of the complex to its final destination (e.g., plasma membrane) (Coleman and Molday, 2011; Shin and Takatsu, 2019). At the plasma membrane, the complex mediates selective phospholipid flipping to maintain the asymmetry of the plasma membrane interior and exterior leaflets (Best et al., 2019; Hankins et al., 2015). For example, these flippase complexes mediate the flipping of anionic phospholipids (e.g., phosphatidylserine (PS)) from the outer to inner leaflet of the phospholipid bilayer (Hankins et al., 2015).

To examine the role of TMEM30A in membrane asymmetry in K562 cells, we ascertained the relative amount of PS in the outer leaflet via a flow cytometric assay using PS-binding Annexin V conjugated to FITC. TMEM30AKO cell lines showed a drastic increase in Annexin V staining, with >90% of the cells being Annexin V positive and the population exhibiting an ∼6-fold increase in Annexin V staining (**Figure 4A, B**). Dead cells were removed from this analysis by staining with SYTOX-Deep Red, ensuring that the Annexin V staining is not due to dead cells. To directly measure PS flippase activity, we incubated cells with fluorescently-tagged nitrobenzoxadiazole (NBD) PS and then used BSA to remove any NBD-PS remaining in the outer PM leaflet. The fluorescence–which represents PS that has flipped into the cytoplasmic leaflet–can then be measured by flow cytometry. As expected, TMEM30AKO cells exhibited a significant impairment in PS flipping (**Figure 4C**).

**Figure 4.**
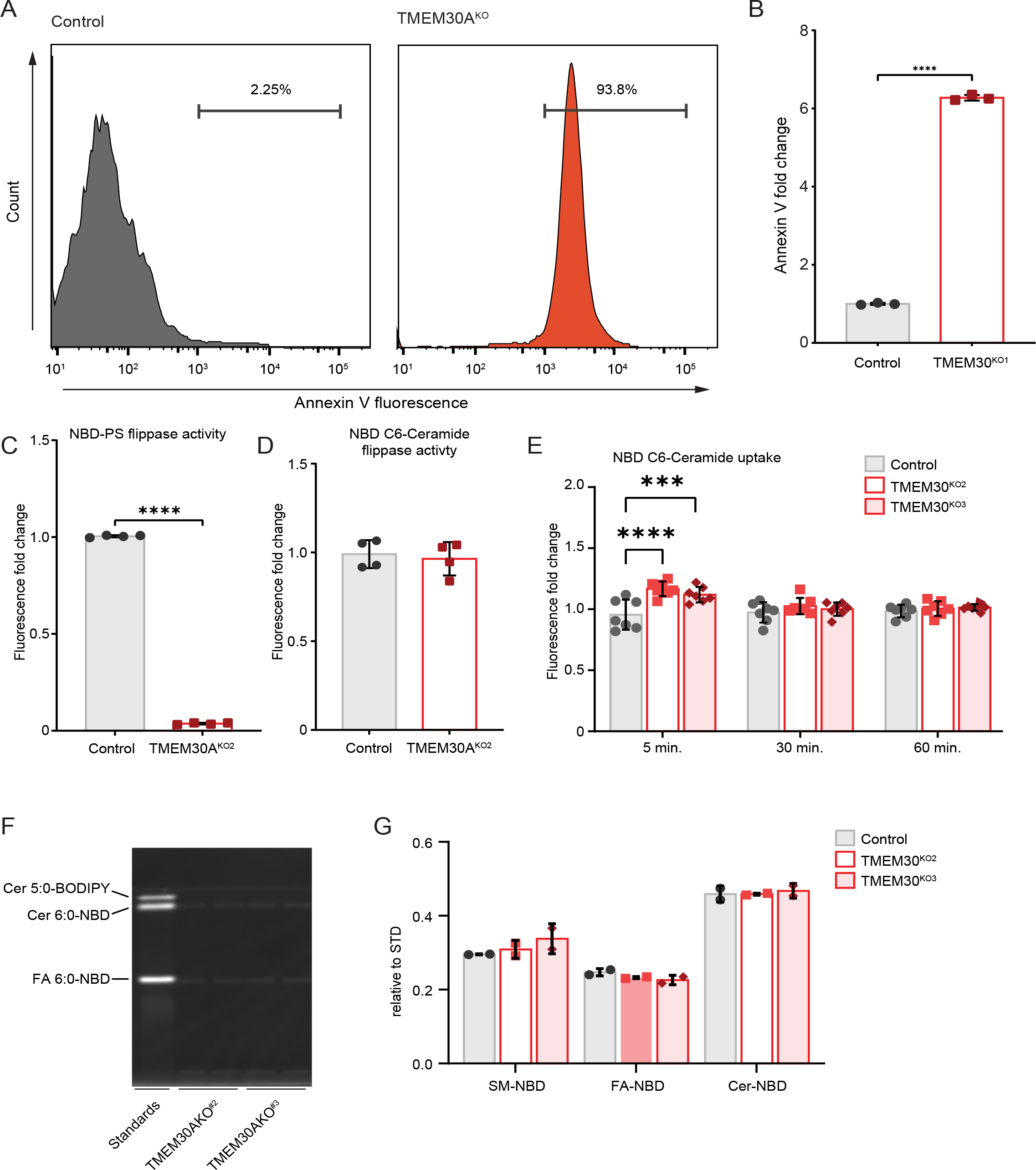
Loss of TMEM30A disrupts phosphatidylserine flipping but not ceramide flipping nor uptake. A) Flow cytometry histograms of annexin V fluorescence in control and TMEM30A^KO1^ cells. B) Quantification of annexin V fluorescence by flow cytometry (as in panel A). C) Quantification of NBD-PS flippase activity by flow cytometry. D) Quantification of NBD C6-Cer flippase activity by flow cytometry. E,F,G) Quantification of NBD C6-Cer uptake by flow cytometry (E) and thin layer chromatography (F,G).

Ceramide is thought to spontaneously flip between bilayer membrane leaflets, an action independent of a protein-based transporter or flippase. We considered the possibility that the altered composition of the TMEM30AKO inner and outer plasma membrane leaflets could influence C6-Cer flipping and cellular uptake. Using NBD C6-Cer, we performed a similar cell-based lipid flipping assay. In contrast to NBD-PS (**Figure 4C**), NBD C6-Cer flipping was unaffected in the TMEM30AKO cells (**Figure 4D**). We also examined longer incubation times to determine if NBD C6-Cer uptake into cells is affected. Although there was a small, statistically significant increase in NBD C6-Cer fluorescence in the TMEM30AKO cells at 5 min, there was no difference at 30 or 60 min (**Figure 4E**). To validate our ceramide uptake assay, we also measured NBD C6-Cer uptake using thin layer chromatography. In addition to the NBD C6-Cer, we observed a band corresponding to the NBD-labelled fatty acid, indicating metabolism of the NBD C6-Cer following cellular uptake (**Figure 4F,G**). Loss of TMEM30A had no effect on the levels of NBD C6-Cer or the NBD-conjugated fatty acid (**Figure 4F,G**). Together, our data indicate that although TMEM30AKO cells exhibit reduced PS flipping and altered membrane asymmetry, there is no defect in C6-Cer flipping or uptake.

### TMEM30A trafficking of ATP11B to the plasma membrane impacts C6-Cer toxicity

The canonical role of TMEM30A is as a subunit of phospholipid flippase complexes that is required both for the trafficking of the flippase complex to the plasma membrane. Indeed, we observed alterations in TMEM30AKO cells PS flippase activity and plasma membrane composition (**Figure 4**). These results raise the possibility that the alteration in plasma membrane asymmetry is responsible for the C6-Cer resistance of TMEM30AKO cells. No P4-type ATPases were identified in the 1% FDR cut-off of our genetic screen, possibly reflecting compensation due to the overlapping functions of these proteins. To identify P4-type ATPases that exhibit impaired trafficking in the TMEM30AKO cells, we implemented a proteomics workflow to examine changes in the surface proteome (**Figure 5A**). 3442 proteins were identified, including many expected plasma membrane proteins (**Supplementary Table S2)**. As anticipated, TMEM30A was reduced in the TMEM30A^KO^ cell lines (**Figure 5B)**. ATP11B and ATP11C, two P4-type ATPase that associate with TMEM30A, exhibited reduced plasma membrane abundance in multiple TMEM30A^KO^ cell lines (**Figure 5B)**. We focused on ATP11B because of its higher abundance, based upon the number of identified spectral counts (**Supplementary Table S2**). The reduction in ATP11B levels was not due to altered gene expression in the TMEM30A^KO^ cells (**Figure 5C**).

**Figure 5.**
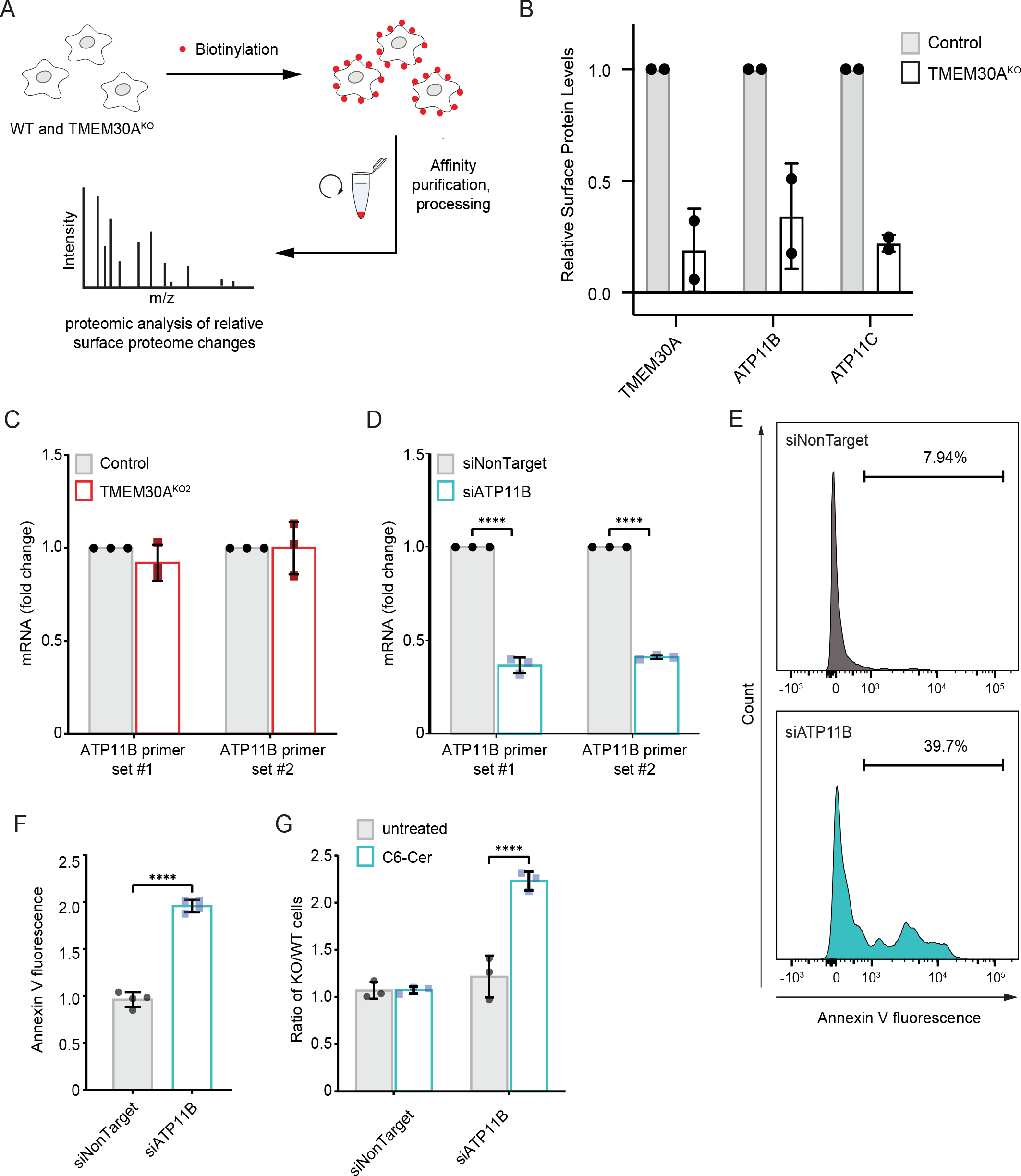
Impaired plasma membrane trafficking of ATP11B impacts PS flipping and exogenous C6-Cer toxicity. A) Schematic of surface proteomics approach. B) Quantification of the relative surface proteome levels for TMEM30A, ATP11B, and ATP11C in control and TMEM30A^KO2^ cells. C) Quantification of ATP11B mRNA by RT-PCR in control and TMEM30A^KO2^ cells. D) Quantification of ATP11B mRNA by RT-PCR in cells transfected with siRNAs, siNonTarget and siATP11B. E) Flow cytometry histograms of annexin V fluorescence following transfection with siNonTarget and siATP11B. F) Quantification of annexin V fluorescence following transfection with siNonTarget and siATP11B (as in panel D). G) C6-Cer competitive growth assay of cells transfected with the indicated siRNA against an untransfected control cell line.

To examine the functional role of ATP11B, we depleted ATP11B using siRNAs (**Figure 5D**). ATP11B depletion resulted in an increase in the percentage of Annexin V positive cells and fluorescence (**Figure 5E,F**), consistent with a role for ATP11B in the maintenance of membrane asymmetry in K562 cells. Similar to the loss of TMEM30A, the depletion of ATP11B also resulted in resistance to C6-Cer toxicity (**Figure 5G**). The effects of ATP11B on membrane asymmetry and C6-Cer resistance were not as large as in the TMEM30AKO cells, possibly due to the partial depletion of ATP11B or the contribution of additional P4-type ATPases such as ATP11C which was also reduced in the TMEM30A^KO^ cells. Together, these results suggest that TMEM30A influences C6-Cer resistance, at least in part, by promoting ATP11B trafficking and plasma membrane asymmetry.

## DISCUSSION

Ceramide plays established roles in apoptosis through its regulation of key signalling events (Morad and Cabot, 2013). The mechanisms by which cancer cells evade ceramide-induced cell death remain incompletely understood. We employed a genome wide CRISPR-Cas9 screen to provide a resource of high confidence genetic modifiers of C6-Cer induced cell death in K562 cells and to reveal a role for plasma membrane symmetry in governing the sensitivity of cells to C6-Cer toxicity.

Our genetic screen identified a host of genes from diverse functional categories; these included enrichments in ceramide and sphingolipid metabolism, membrane biology, vesicular trafficking, and transcription. We provide initial validation of several candidate regulators, including CERS2, GBA, UGCG, CHD8, ARL5, and TMEM30A. Further studies will be required to understand the mechanisms that mediate their role in C6-Cer induced cell death. It is notable that we identified the TNF receptor TNFRSF1A (also known as TNFR1) and NSMAF in our screen using C6-Cer. In the extrinsic apoptotic pathway, TNF binding to TNF receptors leads to ceramide generation whilst NSMAF couples the TNF receptor to neutral sphingomyelinase to produce ceramide from sphingomyelin catabolism. The identification of TNFRSF1A suggests that our screen may detect factors that will influence endogenous ceramide apoptotic signalling in addition to the cell death triggered by the more artificial exogenous C6-Cer treatment.

One of the most interesting discoveries was that the loss of TMEM30A results in resistance to C6-Cer toxicity. TMEM30A is a subunit in phospholipid flippase complexes (Hankins et al., 2015). It binds several P4-type ATPases and is integral to proper ATPase trafficking and function (Coleman and Molday, 2011). Our proteomic analyses of the surface proteome in TMEM30AKO cells revealed a loss of ATP11B, reaffirming the requirement of TMEM30A for trafficking of P4-type ATPases. RNAi-mediated depletion of ATP11B phenocopied the loss of TMEM30A, resulting in the localization of PS to the outer leaflet of the plasma membrane and in resistance to C6-Cer toxicity. This result supports the model that loss of plasma membrane asymmetry is responsible for the C6-Cer resistance phenotype in the TMEM30AKO cells. The effects on PS exposure and C6-Cer resistance were more modest in the ATP11B depleted cells than in the TMEM30AKO cells and it remains possible that other P4-type ATPases also contribute to the observed phenotypes. Ceramide is known to spontaneously flip within the bilayer independent of a protein-based transporter. We did not observe any defect in C6-Cer flipping or uptake, indicating that the resistance phenotype is not simply due to a reduction in C6-Cer access to the interior of the cell.

The plasma membrane is a complex combination of phospholipids, sterols, glycolipids, and proteins (Harayama and Riezman, 2018). In addition, the composition of the inner (cytoplasmic) and outer (exoplasmic) leaflets of the plasma membrane differ (Doktorova et al., 2020). Employing energy generated from ATP hydrolysis, lipid flippases transfer lipids from the outer leaflet to the inner leaflet (Hankins et al., 2015). Flippases act together with floppases– which transfer lipids from the inner to outer leaflets–and scramblases–which mediate bidirectional lipid transport–to dynamically control transbilayer lipid compositions (Hankins et al., 2015). Elegant lipidomics studies indicate that the outer leaflet of the plasma membrane is primarily composed of phosphatidylcholine and sphingolipids, which tend to pack tightly and contribute to a more highly ordered and rigid membrane (Doktorova et al., 2020). The inner leaflet of the plasma membrane is enriched in charged lipids, including PS, phosphatidylethanolamine, phosphatidylinositol, and overall contains more highly unsaturated fatty acids (Doktorova et al., 2020). The canonical enrichment of charged lipids within the inner leaflet contributes to the electrostatic potential across the bilayer (Hankins et al., 2015; Harayama and Riezman, 2018). This influences the insertion and folding of integral membrane proteins as well as the association of peripheral proteins through polybasic stretches (Harayama and Riezman, 2018). As anticipated, we found that PS aberrantly localizes to the outer leaflet of the plasma membrane in the TMEM30AKO cells. The transbilayer distribution of other lipids, such as sphingolipids, in the TMEM30AKO is not known but broader alterations in the composition of plasma membrane inner and outer leaflets may contribute to our observed phenotypes.

While our results implicate plasma membrane asymmetry in cellular sensitivity to C6-Cer toxicity, the exact mechanism is not clear and will require additional studies. Alterations in membrane asymmetry may influence sphingolipid and ceramide metabolic pathways, which would result in changes in intracellular ceramide levels that would affect the formation rate of ceramide-enriched platforms and death-induced signaling complexes. In addition, the altered plasma membrane asymmetry likely influences integral membrane protein topologies and charge-based recruitment of peripheral proteins to the plasma membrane. Plasma membrane asymmetry may also influence the ability of ceramide to form channels that permeabilize membranes, though this activity of ceramide is controversial.

Together, our study provides a global overview of the genetic landscape that governs C6-Cer toxicity. Moreover, our findings implicate plasma membrane asymmetry as a key factor in the cellular sensitivity to C6-Cer, setting the stage for future studies examining the connection between specific plasma membrane lipids and ceramide metabolism, plasma membrane physical properties, and cell death pathways.

## METHODS

### Cell lines and culture conditions

HEK293T was obtained from UC Berkeley’s cell culture facility. K562 and K562-Cas9-BFP cells were a generous donation from R. Kopito (Stanford University). HEK293T was maintained in DMEM containing 4.5 g/l glucose and L-glutamine (Corning, 10-017). K562 cells and their derivatives were maintained in RPMI-1640 with L-glutamine (Corning, 10-040-CM). All media was supplemented with 10% fetal bovine serum (Gemini Bio Products) and penicillin (10,000 I.U mL^-1^). Cell lines were maintained at 37°C with 5% CO_2_ with regular screening for mycoplasma contamination.

### EC_50_ Death Assay

Cell death was assayed via the Essen IncuCyte Live-Cell imaging system (Sartorius). Ten thousand K562 cells per well were seeded in black 96-well plates (Corning, 3904). Media containing a final concentration of 15nM SYTOX-Green Dead Cell Stain (Invitrogen, S34860) and C6-Ceramide (d18:1/6:0) (Enzo, BML-SL110) at various concentrations was added to produce a final cell density of 500,000 cells/mL. Plates were carefully transferred to the IncuCyte system (kept at 37°C with 5% CO_2_) and imaged for 24hours (24hr). Three images per well were captured in the green and brightfield channels every hour for the treatment period. The Sartorius image analysis software outputs the number of green objects (SYTOX-Green positive, i.e., dead cells) as well as the total number of objects (brightfield segmentation). For each C6-Cer concentration, the ratio of dead objects over total objects was plotted over the 24hr imaging cycle. From this, Prism (Graphpad) was used to calculate the area under the curve (AUC), plotted as function of C6-Cer dosage and mathematically determine the EC_50_.

### CRISPR-Cas9 synthetic lethal screen

The screen was performed as previously described^1^. In brief, the Bassik Human CRISPR Knockout Library (Addgene, 101926-101934) is separated into 9 sublibraries comprising a total of 225,171 elements and targeting approximately 20,500 genes (10 sgRNAs per target). To generate the lentiviral library pool, each sublibrary pool was co-transfected with third-generation lentiviral packaging plasmids (pVSV-G/MD2.G [Addgene, 12259], pRSV-Rev [Addgene, 12253] and pMDLg [Addgene, 12251]) into low-passage HEK293T cells. Lentiviral-containing media was collected at 48hr and at 72hr post-transfection. These media pools were then combined and filtered via 0.45-micron cartridge. The resultant lentiviral media was used to infect approximately 2.0 × 10^8^ K562-Cas9 cells for 72hr. After this infection period, the cells were spun down at 500 × g for 5min and viral-laden media was removed. These cells then underwent 5μg/mL Puromycin Dihydrochloride (Gibco, A1113803) selection until the population was over 97% mCherry positive. Cells then recovered in puromycin-free media until a total of 2.5 × 10^8^ (∼1,000-fold total library elements). This pool was then split and treated with either ethanol (EtOH) or 30μM C6-Cer for 24hr. The treatment period was followed by a 48hr recovery and subsequently, the treatment cycle was repeated thrice more. Each subsequent cycle required a slight uptick in C6-Cer concentration as the pools experienced rapid resistance to death and thus the drug concentration was raised to achieve ∼50% death. The dosage for cycle 2 and upwards is as follows: 32.5μM, 35μM and finally, 40μM. Throughout the screen, K562 cells are maintained at 500,000 cells/mL.

After the last bout of treatment, cells were twice-washed with PBS, pelleted into numbers approximating 250-fold of total library elements and stored at -80°C. Genomic DNA was extracted using the QIAmp DNA Blood Maxi Kit (Qiagen) according to the manufacturer’s directions. This genomic DNA underwent two rounds of PCR to first amplify sgRNA sequences and then index them using Illumina TruSeq LT adaptor sequences.

PCR amplicons from each sample were pooled into a 1:1 ratio (EtOH:C6-Cer) based on concentrations determined by Qubit Fluorometric Quantification (Invitrogen). Deep sequencing was done on an Illumina NextSeq instrument at the Oklahoma Medical Research Foundation. Sequence reads were trimmed, then aligned using Bowtie with zero mismatches tolerated. Enrichment, confidence scores and p-values were calculated using the casTLE Python code as previously described^2^.

### Generation of CRISPR-Cas9 genome-edited cell lines

Knockout cell lines were generated the same way as the lentiviral portion of the CRISPR-Cas9 screen. Cas9 cutting functionality was validated via infecting K562-Cas9 with a self-cutting mCherry plasmid (a kind gift from M. Bassik, Stanford University) that expresses mCherry as well as a sgRNA sequence targeting said gene.

sgRNA sequences selected from the CRISPR KO screen were used to create the individual transfection plasmids. All guide sequences were inserted into pMCB320 between the BstXI and BlpI sites as previously described^3^.

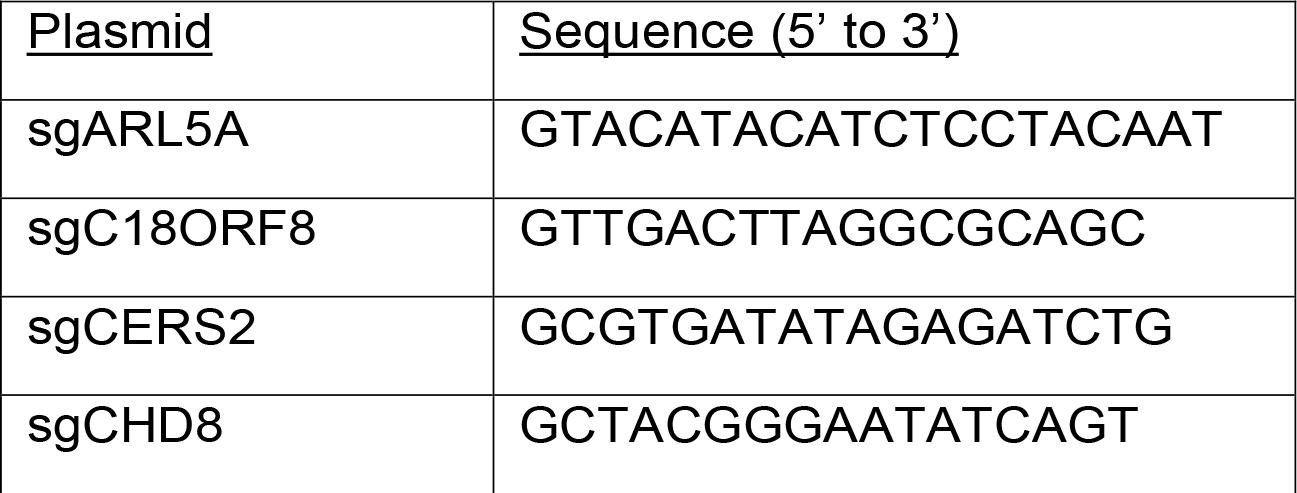
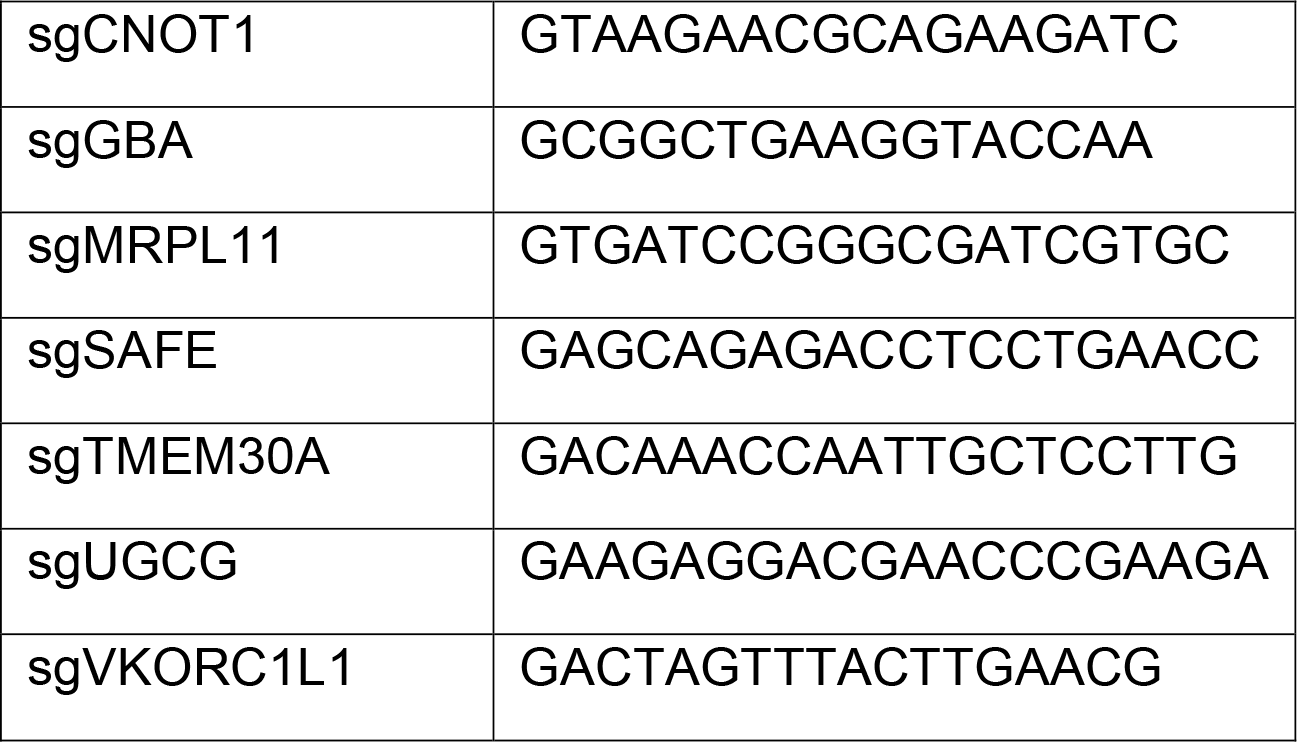

Single clones were isolated from the sgTMEM30A polyclonal line to create TMEM30A^KO1^ and TMEM30A^KO2^ via serial dilution. Serial dilution was also employed to form clonal lines of sgARL5A to create ARL5A^KO1^ and ARL5A^KO2^. The sgRNA sequences to the additional knockout lines are as follows:

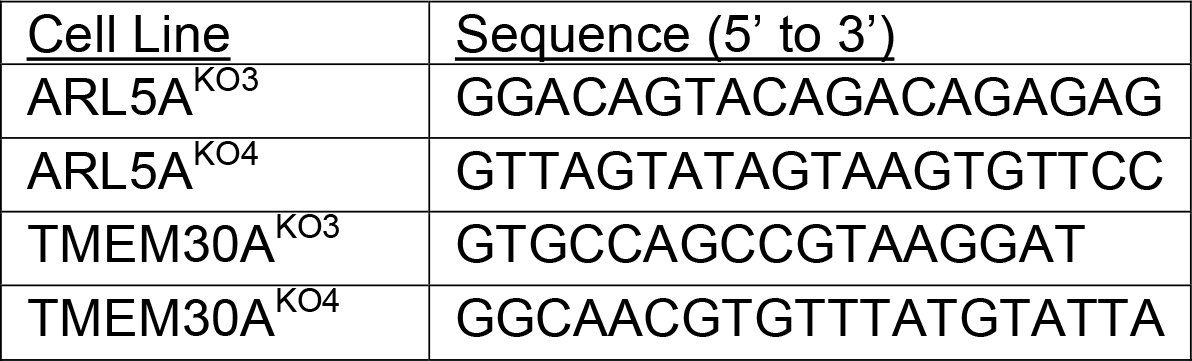

### Competitive Growth Assay

WT cells (self-cut mCherry, no mCherry emission) and KO cells (mCherry positive) were seeded in a 1:1 ratio with a final density of 500,000 cells/mL into a 96-well round-bottom plate. Wells were treated with either vehicle (EtOH, untreated) or C6-Ceramide for 24hr. Cells were then spun down, media aspirated and then replenished with media containing 15nM SYTOX-Green. The plate was assayed via flow cytometry on a BD LSRFortessa in the UC Berkeley Flow Cytometry Facility; dead cells were gated out (positive SYTOX-Green) and live cells were analysed to discover the percentages of mCherry +/-cells. Two-way ANOVA was calculated on Prism.

### Annexin V Assay

Assay was adapted from manufacturer directions of the FITC Annexin V Apoptosis Detection Kit (BD Pharmingen, 556547). In brief, K562 cells were collected from cultures that were 98% live (determined by Trypan Blue staining). Samples were incubated with SYTOX-Deep Red Live/Dead Fluorescent Dye (Invitrogen, S11380) for 15min at 37°C, then washed twice with DPBS (Gibco, 14190-144) before proceeding with Annexin V-FITC incubation. Samples were analysed by flow cytometry; dead cells (i.e., SYTOX-Deep Red positive) were gated out and the FITC spectra measured of the remaining population.

### Flippase Assay

NBD C6-Ceramide (Cayman, 62527) and NBD PS (Avanti, 810194C) were prepared as previously described (Kay et al., 2012). In brief, compounds were dried under N2 and resuspended in methanol. Incorporation of lipids is modified from methods described (Goñi et al., 2014; Sandhoff et al., 2018). Compounds are pre-incubated in 4mg/mL BSA (Sigma, A8806) in H_2_O under constant rotation at 300rpm for 30min at 37°C. K562 cells were collected via centrifugation, washed and equilibrated at 15°C for 30min in Hank’s balanced salt solution (HBSS) containing 1g/L glucose (Gibco, 14025-092) at a concentration of 500,000 cells/mL. After cell equilibration, the incubated compounds were added to the cell suspension to the required lipid concentration. This mixture was incubated at 15°C for 20min with constant rotational speed of 300rpm. At the end of the timepoint, NBD conjugated compounds were mixed with equal volume of ice-cold 5mg/mL BSA in HBSS to extract the lipids incorporated into the exoplasmic PM leaflet. Cells were then analysed by flow cytometry to find the geometric mean of FITC emission. Replicates were averaged and normalised to control values.

### Uptake Assay

Uptake assay is adapted from the flippase method. Samples were plated in a 96-well round bottom plate to a final concentration of 500,000 cells/mL with lipid incorporation occurring in regular media. Samples were incubated at 37°C at 5% CO_2_ for the designated timepoints. At the end of each timepoint, cells were washed thrice with DPBS and assayed by flow cytometry. Replicates were averaged and normalised to control values.

### High Performance Thin-Layer Chromatography

Cells were isolated and washed 2x with PBS. Cells were then pelleted at 500xg for 5 min, the supernatant was removed, and cell pellets were stored at -80°C. Before extraction, cell pellets were thawed on ice for 30 min and resuspended in 50 ul PBS. Lipids were extracted by adding 1250 uL tert-butyl methyl ether (HPLC grade, Sigma-Aldrich, USA) and 375 uL methanol (HPLC grade, Fisher Scientific, USA), both containing 0.1% (w/v) 2,6-Di-tert-butyl-4-methylphenol (GC grade, Sigma-Aldrich, USA). The mixture was incubated on an orbital mixer for 1 h (room temperature, 32 rpm). To induce phase separation, 315 uL water containing 0.1% (w/v) 2,6-Di-tert-butyl-4-methylphenol (GC grade, Sigma-Aldrich, USA) was added, and the mixture was incubated on an orbital mixer for 10 min (room temperature, 32 rpm). Samples were centrifuged (room temperature, 10 min, 17,000 x g). Upper organic phase was collected and subsequently dried in vacuo (Eppendorf concentrator 5301, 1 ppm).

The lipid extract was dissolved in 50 uL chloroform/methanol (2:1, v/v) and 5 ul of each extract were loaded on silica coated, glass-bottomed plates (HPTLC silica gel 60, 10×10 cm, Merck). Pure standards of FA 6:0-NBD (Cayman Chemical, USA), Cer 18:1;O2/6:0-NBD (Cayman Chemical, USA) and Cer 18:1;O2/5:0-BODIPY (Cayman Chemical, USA) were loaded as 0.1 nmol aliquots on the same plates. Lipids were separated using a solvent mixture of triethylamine/chloroform/ethanol/water (5:5:5:1, v/v) (all solvents HPLC grade, Sigma-Aldrich, USA) as mobile phase in a solvent vapour saturated twin trough chamber (CAMAG, Switzerland). Separated lipids were imaged directly on glass-backed TLC plates using a Gel Doc EZ System (BioRad, USA) in combination with the UV Tray filter (BioRad, USA).

Densitometric quantification of lanes was performed using Image Lab Software Version 5.2.1 (BioRad, USA)

### Surface Biotinylation Assay and Proteomics

All buffers were filter sterilised prior to use. Each sample required 24 million cells that were pelted and washed with PBS, twice. 10mM of freshly prepared EZ-Link™ Sulfo-NHS-SS-Biotin (Thermo, A39258) in PBS and added to the cell mixture at 80μL of 10mM NHS-SS-Biotin per mL of sample volume. Samples were incubated by rocking at room temperature for 30min. After incubation, cells were pelleted at 300 g-force for 3 min, excess biotin discarded and then washed twice with ice-cold TBS.

Proteins were collected from cells via lysing with dissolved Pierce™ Protease Inhibitor Mini Tablets, EDTA-free (Thermo, A32955) in RIPA buffer (Thermo, 89901). Samples were sonicated for 15sec at 15% amplitude, followed by vortexing for 10sec every 10min for 30min. This was followed by a 5min centrifugation at 15,000 g-force and supernatant.

Enrichment of biotinylated proteins proceeded via an overnight incubation at 4°C on NeutrAvidin resin (Thermo, 53150) at a ratio of 10μL of beads per 100μg protein. After incubation, samples were centrifuged at 700 g-force for 2min at 4°C. Pellets then underwent a series of wash steps: twice with lysis buffer, thrice with ultrapure H_2_O and finally, twice with 50mM ammonium bicarbonate (ABC), pH 8.0. A portion of clarified proteins were assessed for biotinylation efficiency via western-blotting with a streptavidin secondary antibody (Li-Cor, 32230). Biotinylated proteins were eluted from resin using 10mM dithiothreitol (Thermo, R0861) dissolved in 50mM ABC by end-over-end rotation at room temperature for 30min. Sample was centrifuged at 700 g-force for 2 min and supernatant collected.

Sample preparation for mass spectrometry required the addition of 25 μL of freshly-prepared 55mM iodoacetamide (dissolved in 50mM ABC) with a requisite 30min incubation at room temperature in the dark. After the time interval, we added ice-cold acetone to mix overnight at -20°C. Samples were then centrifuged at 15,000 g-force for 10min, then decanted for 30min at room temperature to precipitate proteins.

Precipitated proteins were resuspended in 25mM ammonium bicarbonate, digested overnight using 1μg of trypsin (Promega, V5113) at 37°C. Samples were acidified with 10% v/v of trifluoroacetic acid, desalted using C18 stage tips, and dried. For MS analysis, peptides were resuspended in 1% formic acid and separated on an Easy nLC 1000 UHPLC equipped with a 15 cm nanoLC column. Using a flow rate of 300 nL/min, the linear gradient was 5% to 35% over B for 90 min, 35% to 95% over B for 5 min, and 95% hold over B for 15 min (solvent A: 0.1% formic acid (FA) in water, solvent B: 0.1% FA in ACN). The table indicates ley mass spectrometer parameters.

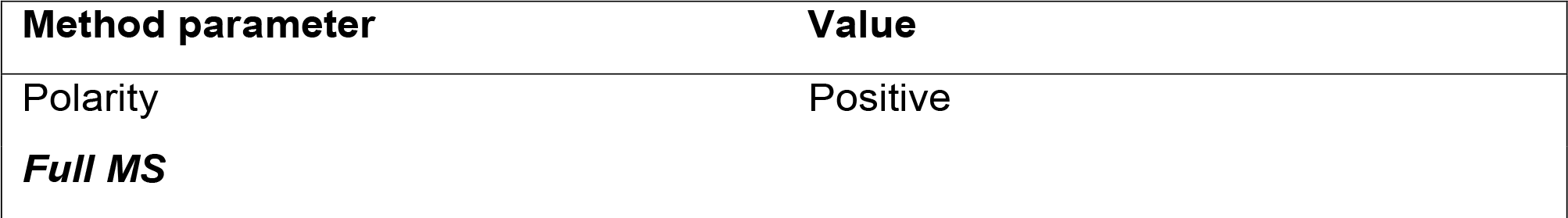
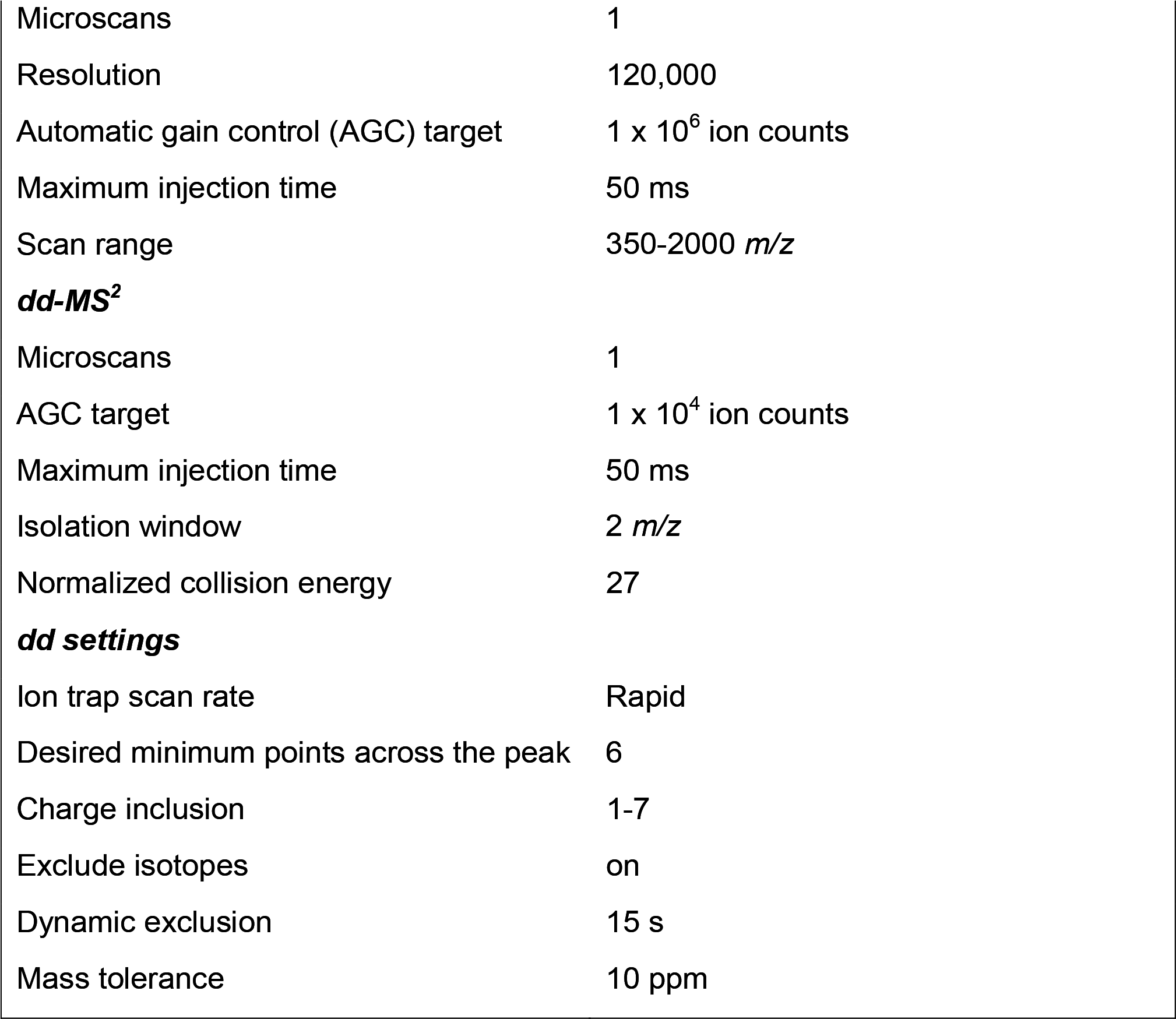

| Method parameter | Value |
| --- | --- |
| Polarity | Positive |
| *Full MS* |  |
| Microscans | 1 |
| Resolution | 120,000 |
| Automatic gain control (AGC) target | 1 x 10<sup>6</sup> ion counts |
| Maximum injection time | 50 ms |
| Scan range | 350-2000 *m/z* |
| ***dd-MS<sup>2</sup>*** |  |
| Microscans | 1 |
| AGC target | 1 x 10<sup>4</sup> ion counts |
| Maximum injection time | 50 ms |
| Isolation window | 2 *m/z* |
| Normalized collision energy | 27 |
| ***dd settings*** |  |
| Ion trap scan rate | Rapid |
| Desired minimum points across the peak | 6 |
| Charge inclusion | 1-7 |
| Exclude isotopes | on |
| Dynamic exclusion | 15 s |
| Mass tolerance | 10 ppm |

### Method parameter Value

Peptide identities and relative abundances were determined using Proteome Discoverer 2.4. Ion chromatograms were extracted using Xcalibur Qual Browser for each peptide of interest with a mass tolerance of 0.5 Da. We thank Dr. Steve Eyles (UMass Amherst, RRID: SCR_019063) for assistance with high-resolution MS acquired on an Orbitrap Fusion mass spectrometer (NIH grant: 1S10OD010645-01A1).

### siRNA Knockdown of ATP11B

sgSAFE cells were transfected with either siATP11B (Horizon, L-023660-00-0005) or siNon-targeting control (Horizon, D-001810-10-05) using Lipofectamine RNAiMax transfection reagent (Thermo, 13778) according to manufacturer directions. Incubation proceeded for 48 hr, whereupon samples were washed and divvied into 3 populations: two for subsequent competitive growth assays and Annexin V studies, with the last assessed for knockdown efficiency via RT-qPCR. K562 self-cutting mCherry cells were also transfected with siNon-targeting control to serve as the mCherry-population in our competitive growth assay.

### RT-qPCR

Total RNA of cell samples was extracted using the Monarch Total RNA kit (NEB, T2010S) and then reverse transcribed with iScript cDNA Synthesis kit (Biorad, 1708890). cDNA was measured using a CFX96 Touch Deep Well Real-Time PCR system (Biorad) with the probe-based PrimeTime Gene Expression Master Mix (IDT, 1055770). Fold changes in cDNA were determined using the ΔΔC_t_ method^8^ normalised to PPIA cDNA. Primer pairs were purchased from IDT’s predesigned PrimeTime Standard qPCR assay; sequences are as follows:

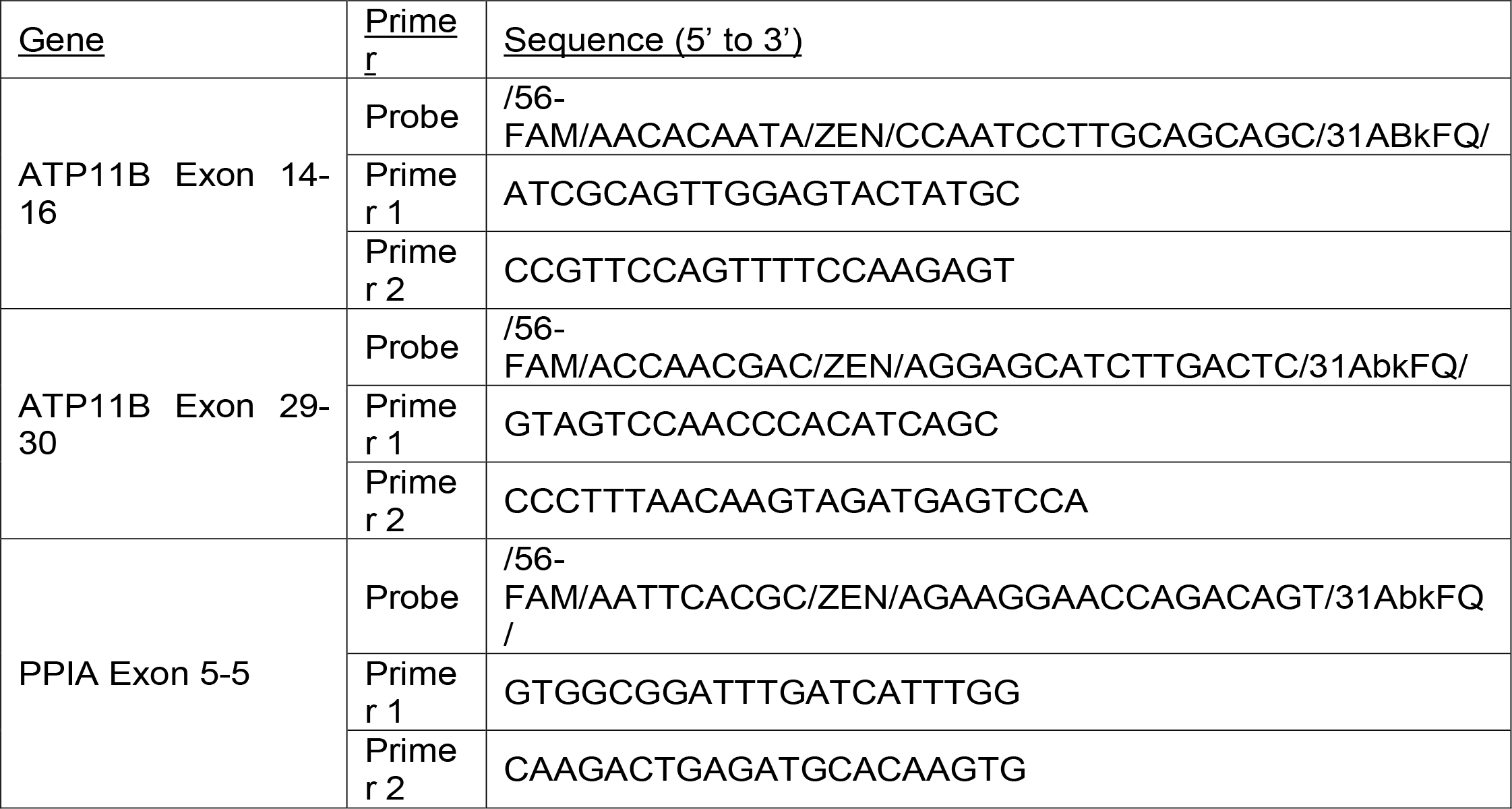

### Cell counts, Statistical Analysis and Reproducibility

Unless otherwise indicated, cell amount/concentrations were derived from live cell counts (determined by Trypan Blue staining). Experimental populations were collected from growing pools that exhibited ≥ 97% live. Additionally, unless otherwise indicated, experiments were done in triplicate with statistical significance calculated in Prism using a two-tailed unpaired t-test (p-value ≤0.0332 [*], ≤0.0021 [**], ≤0.0002 [***], <0.0001 [****]).

## ACKNOWLEDGEMENTS

This research was supported by grants from the National Institutes of Health (R01GM112948 to J.A.O. and F31GM134645 to S.N.S.M.) and American Cancer Society (Research Scholar Award RSG-19-192-01 to J.A.O.). J.A.O. is a Chan Zuckerberg Biohub investigator. We thank Dr. Steve Eyles (UMass Amherst, RRID: SCR_019063) for assistance with high-resolution MS acquired on an Orbitrap Fusion mass spectrometer funded by NIH grant 1S10OD010645-01A1.

## AUTHOR CONTRIBUTIONS

S.N.S.M. and J.A.O. conceived of the project, designed the experiments, and wrote the majority of the manuscript. All authors read, edited, and contributed to the manuscript. S.N.S.M. and K.K.D. performed the proteomics experiments and S.N.S.M and M.L. performed the thin layer chromatography experiments.

## AUTHOR INFORMATION

Correspondence and requests for materials should be addressed to J.A.O.

## SUPPLEMENTARY FIGURE LEGENDS

**Figure S1.**
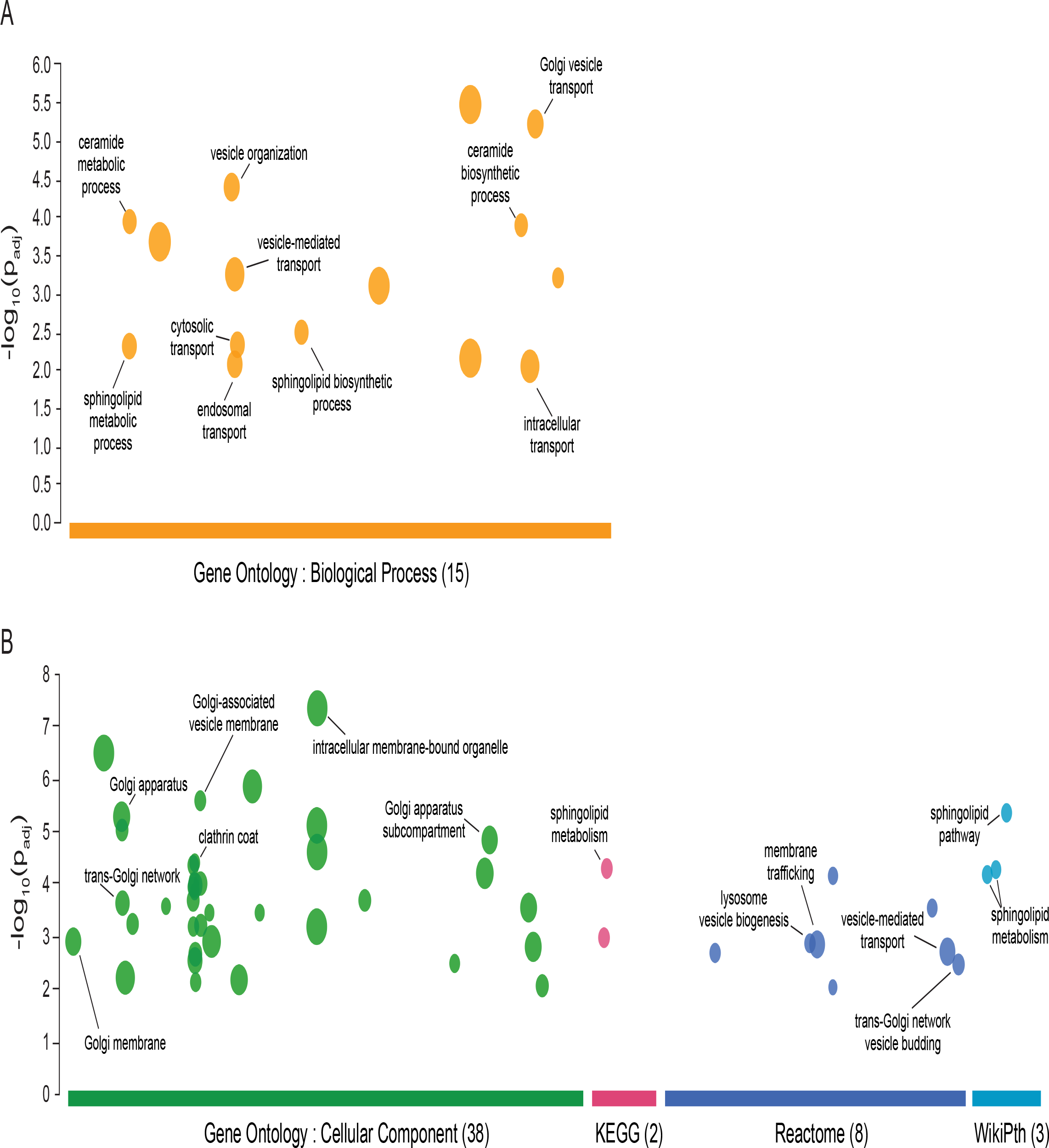
Analysis of C6-Cer genetic modifiers. Significantly enriched and disenriched genes from the genome-wide C6-Cer CRISPR screen were analyzed using g:Profiler to identify functional relationships.

**Figure S2.**
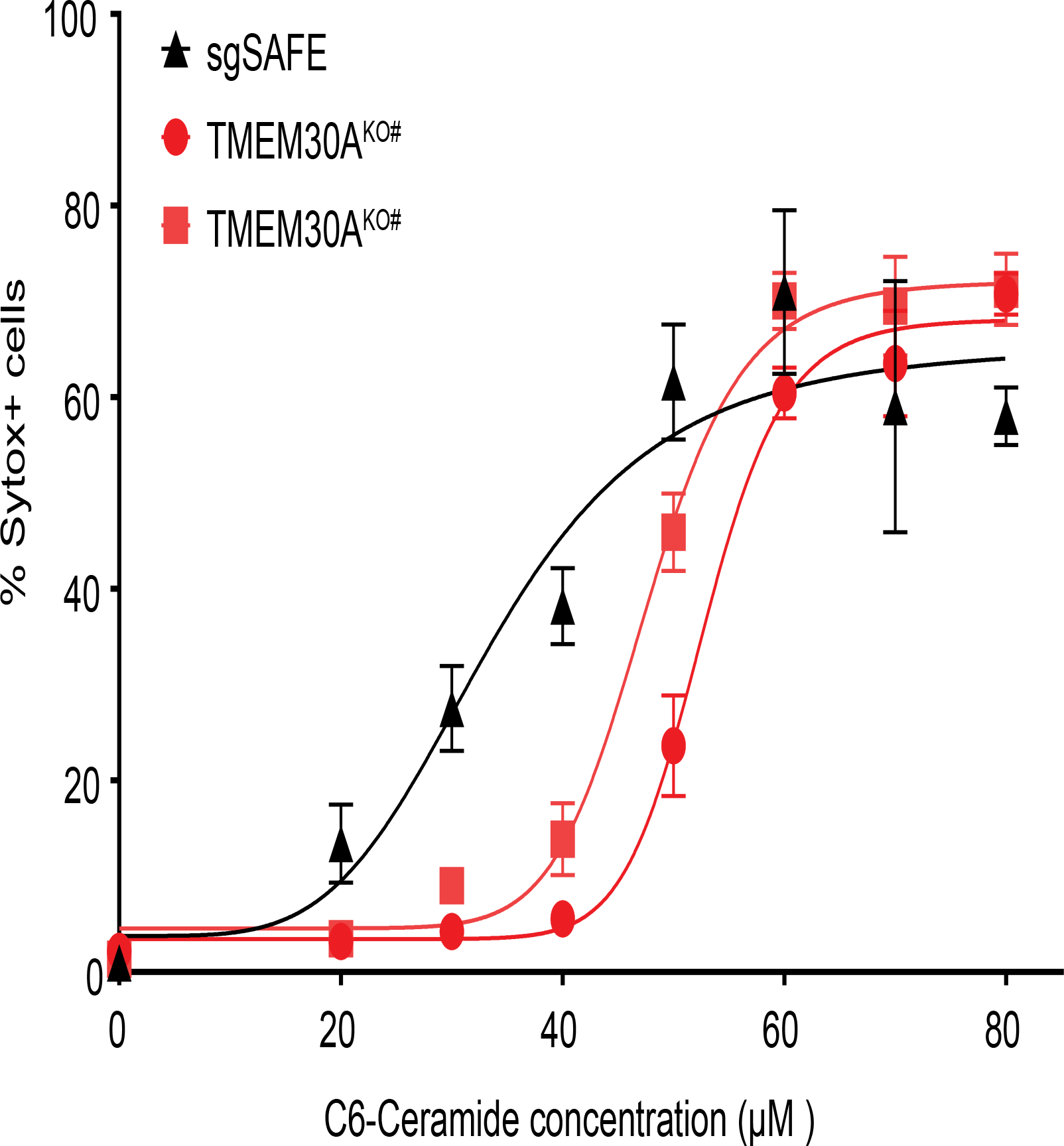
Loss of TMEM30A sensitizes cells to exogenous C6-Cer-induced cell death. K562 cells were treated with the indicated concentrations of C6-Cer for 24 hr and the percentage of dead cells (Sytox+) quantified using flow cytometry.

## SUPPLEMENTARY TABLES

**Supplementary Table S1**.

CRISPR-Cas9 screen data.

**Supplementary Table S2**.

Surface proteome analysis.

## REFERENCES

1. Morad, S. A. F. & Cabot, M. C. Ceramide-orchestrated signalling in cancer cells. Nat. Rev. Cancer 13, 51–65 (2013).

2. Moro, K., Nagahashi, M., Gabriel, E., Takabe, K. & Wakai, T. Clinical application of ceramide in cancer treatment. Breast Cancer 26, 407–415 (2019).

3. Park, J.-W., Park, W.-J. & Futerman, A. H. Ceramide synthases as potential targets for therapeutic intervention in human diseases. Biochim. Biophys. Acta 1841, 671–681 (2014).

4. Oskouian, B. & Saba, J. D. Cancer treatment strategies targeting sphingolipid metabolism. Adv. Exp. Med. Biol. 688, 185–205 (2010).

5. Sandhoff, R., Schulze, H. & Sandhoff, K. Ganglioside metabolism in health and disease. Prog Mol Biol Transl Sci 156, 1–62 (2018).

6. Goñi, F. M., Sot, J. & Alonso, A. Biophysical properties of sphingosine, ceramides and other simple sphingolipids. Biochem. Soc. Trans. 42, 1401–1408 (2014).

7. Goñi, F. M. & Alonso, A. Biophysics of sphingolipids I. Membrane properties of sphingosine, ceramides and other simple sphingolipids. Biochim. Biophys. Acta 1758, 1902–1921 (2006).

8. Hannun, Y. A. & Obeid, L. M. Principles of bioactive lipid signalling: lessons from sphingolipids. Nat. Rev. Mol. Cell Biol. 9, 139–150 (2008).

9. Pewzner-Jung, Y., Ben-Dor, S. & Futerman, A. H. When do Lasses (longevity assurance genes) become CerS (ceramide synthases)?: Insights into the regulation of ceramide synthesis. J. Biol. Chem. 281, 25001–25005 (2006).

10. Hanada, K., Kumagai, K., Tomishige, N. & Kawano, M. CERT and intracellular trafficking of ceramide. Biochim. Biophys. Acta 1771, 644–653 (2007).

11. Gómez-Muñoz, A. Ceramide 1-phosphate/ceramide, a switch between life and death. Biochim. Biophys. Acta 1758, 2049–2056 (2006).

12. Kitatani, K., Idkowiak-Baldys, J. & Hannun, Y. A. The sphingolipid salvage pathway in ceramide metabolism and signaling. Cell Signal. 20, 1010–1018 (2008).

13. Castro, B. M., Prieto, M. & Silva, L. C. Ceramide: a simple sphingolipid with unique biophysical properties. Prog. Lipid Res. 54, 53–67 (2014).

14. Nica, A. F. et al. Ceramide promotes apoptosis in chronic myelogenous leukemia-derived K562 cells by a mechanism involving caspase-8 and JNK. Cell Cycle 7, 3362–3370 (2008).

15. Gencer, E. B., Ural, A. U., Avcu, F. & Baran, Y. A novel mechanism of dasatinib-induced apoptosis in chronic myeloid leukemia; ceramide synthase and ceramide clearance genes. Ann. Hematol. 90, 1265–1275 (2011).

16. Yagci, Z. B., Esvap, E., Ozkara, H. A., Ulgen, K. O. & Olmez, E. O. Inflammatory response and its relation to sphingolipid metabolism proteins: Chaperones as potential indirect anti-inflammatory agents. Adv. Protein Chem. Struct. Biol. 114, 153–219 (2019).

17. Fillet, M. et al. Mechanisms involved in exogenous C2- and C6-Cer-induced cancer cell toxicity. Biochem. Pharmacol. 65, 1633–1642 (2003).

18. Voelkel-Johnson, C., Hannun, Y. A. & El-Zawahry, A. Resistance to TRAIL is associated with defects in ceramide signaling that can be overcome by exogenous C6-Cer without requiring down-regulation of cellular FLICE inhibitory protein. Mol. Cancer Ther. 4, 1320– 1327 (2005).

19. Ji, C. et al. Exogenous cell-permeable C6-Cer sensitizes multiple cancer cell lines to Doxorubicin-induced apoptosis by promoting AMPK activation and mTORC1 inhibition. Oncogene 29, 6557–6568 (2010).

20. Takeda, S., Mitsutake, S., Tsuji, K. & Igarashi, Y. Apoptosis occurs via the ceramide recycling pathway in human HaCaT keratinocytes. J. Biochem. 139, 255–262 (2006).

21. Jiang, F. & Doudna, J. A. CRISPR-Cas9 Structures and Mechanisms. Annu. Rev. Biophys. 46, 505–529 (2017).

22. Morgens, D. W., Deans, R. M., Li, A. & Bassik, M. C. Systematic comparison of CRISPR/Cas9 and RNAi screens for essential genes. Nat. Biotechnol. 34, 634–636 (2016).

23. Deans, R. M. et al. Parallel shRNA and CRISPR-Cas9 screens enable antiviral drug target identification. Nat. Chem. Biol. 12, 361–366 (2016).

24. Morgens, D. W. et al. Genome-scale measurement of off-target activity using Cas9 toxicity in high-throughput screens. Nat. Commun. 8, 15178 (2017).

25. Andersson, L. C., Nilsson, K. & Gahmberg, C. G. K562--a human erythroleukemic cell line. Int. J. Cancer 23, 143–147 (1979).

26. Leto, D. E. et al. Genome-wide CRISPR Analysis Identifies Substrate-Specific Conjugation Modules in ER-Associated Degradation. Mol. Cell 73, 377–389.e11 (2019).

27. Benjamini, Y., Krieger, A. M. & Yekutieli, D. Adaptive linear step-up procedures that control the false discovery rate. Biometrika 93, 491–507 (2006).

28. Siskind, L. J. & Colombini, M. The lipids C2- and C16-ceramide form large stable channels. Implications for apoptosis. J. Biol. Chem. 275, 38640–38644 (2000).

29. Levy, M. & Futerman, A. H. Mammalian ceramide synthases. IUBMB Life 62, 347–356 (2010).

30. Laviad, E. L. et al. Characterization of ceramide synthase 2: tissue distribution, substrate specificity, and inhibition by sphingosine 1-phosphate. J. Biol. Chem. 283, 5677–5684 (2008).

31. Ohno, Y. et al. ELOVL1 production of C24 acyl-CoAs is linked to C24 sphingolipid synthesis. Proc. Natl. Acad. Sci. USA 107, 18439–18444 (2010).

32. Sassa, T. et al. Impaired epidermal permeability barrier in mice lacking elovl1, the gene responsible for very-long-chain fatty acid production. Mol. Cell. Biol. 33, 2787–2796 (2013).

33. Sassa, T., Suto, S., Okayasu, Y. & Kihara, A. A shift in sphingolipid composition from C24 to C16 increases susceptibility to apoptosis in HeLa cells. Biochim. Biophys. Acta 1821, 1031–1037 (2012).

34. Spiegel, S. & Milstien, S. Sphingosine-1-phosphate: an enigmatic signalling lipid. Nat. Rev. Mol. Cell Biol. 4, 397–407 (2003).

35. Wegner, M.-S. et al. UDP-glucose ceramide glucosyltransferase activates AKT, promoted proliferation, and doxorubicin resistance in breast cancer cells. Cell Mol. Life Sci. 75, 3393– 3410 (2018).

36. Wegner, M.-S., Gruber, L., Mattjus, P., Geisslinger, G. & Grösch, S. The UDP-glucose ceramide glycosyltransferase (UGCG) and the link to multidrug resistance protein 1 (MDR1). BMC Cancer 18, 153 (2018).

37. Osawa, Y. et al. Roles for C16-ceramide and sphingosine 1-phosphate in regulating hepatocyte apoptosis in response to tumor necrosis factor-alpha. J. Biol. Chem. 280, 27879–27887 (2005).

38. Reimand, J., Kull, M., Peterson, H., Hansen, J. & Vilo, J. g:Profiler--a web-based toolset for functional profiling of gene lists from large-scale experiments. Nucleic Acids Res. 35, W193–200 (2007).

39. Raudvere, U. et al. g:Profiler: a web server for functional enrichment analysis and conversions of gene lists (2019 update). Nucleic Acids Res. 47, W191–W198 (2019).

40. Reimand, J. et al. Pathway enrichment analysis and visualization of omics data using g:Profiler, GSEA, Cytoscape and EnrichmentMap. Nat. Protoc. 14, 482–517 (2019).

41. Kanehisa, M., Sato, Y., Furumichi, M., Morishima, K. & Tanabe, M. New approach for understanding genome variations in KEGG. Nucleic Acids Res. 47, D590–D595 (2019).

42. Jassal, B. et al. The Reactome Pathway Knowledgebase. Nucleic Acids Res. 48, D498– D503 (2020).

43. Slenter, D. N. et al. WikiPathways: a multifaceted pathway database bridging metabolomics to other omics research. Nucleic Acids Res. 46, D661–D667 (2018).

44. Thul, P. J. et al. A subcellular map of the human proteome. Science 356, (2017).

45. Lacombe, J. & Ferron, M. VKORC1L1, an enzyme mediating the effect of vitamin K in liver and extrahepatic tissues. Nutrients 10, (2018).

46. Oldenburg, J., Watzka, M. & Bevans, C. G. VKORC1 and VKORC1L1: Why do Vertebrates Have Two Vitamin K 2,3-Epoxide Reductases? Nutrients 7, 6250–6280 (2015).

47. Casalou, C., Ferreira, A. & Barral, D. C. The role of ARF family proteins and their regulators and effectors in cancer progression: A therapeutic perspective. Front. Cell Dev. Biol. 8, 217 (2020).

48. Wang, Q. et al. microRNA-202-3p inhibits cell proliferation by targeting ADP-ribosylation factor-like 5A in human colorectal carcinoma. Clin. Cancer Res. 20, 1146–1157 (2014).

49. Shi, M. et al. Amino acids stimulate the endosome-to-Golgi trafficking through Ragulator and small GTPase Arl5. Nat. Commun. 9, 4987 (2018).

50. Donaldson, J. G. & Jackson, C. L. ARF family G proteins and their regulators: roles in membrane transport, development and disease. Nat. Rev. Mol. Cell Biol. 12, 362–375 (2011).

51. Ishida, M. & Bonifacino, J. S. ARFRP1 functions upstream of ARL1 and ARL5 to coordinate recruitment of distinct tethering factors to the trans-Golgi network. J. Cell Biol. 218, 3681– 3696 (2019).

52. Yang, F. et al. Deletion of a flippase subunit Tmem30a in hematopoietic cells impairs mouse fetal liver erythropoiesis. Haematologica 104, 1984–1994 (2019).

53. Yang, Y., Liu, W., Sun, K., Jiang, L. & Zhu, X. Tmem30a deficiency leads to retinal rod bipolar cell degeneration. J. Neurochem. 148, 400–412 (2019).

54. Coleman, J. A. & Molday, R. S. Critical role of the beta-subunit CDC50A in the stable expression, assembly, subcellular localization, and lipid transport activity of the P4-ATPase ATP8A2. J. Biol. Chem. 286, 17205–17216 (2011).

55. He, Y., Xu, J., Wu, X. & Li, L. Structures of a P4-ATPase lipid flippase in lipid bilayers. Protein Cell 11, 458–463 (2020).

56. Palmgren, M., Østerberg, J. T., Nintemann, S. J., Poulsen, L. R. & López-Marqués, R. L. Evolution and a revised nomenclature of P4 ATPases, a eukaryotic family of lipid flippases. Biochim. Biophys. Acta Biomembr. 1861, 1135–1151 (2019).

57. Nishiyama, M. et al. CHD8 suppresses p53-mediated apoptosis through histone H1 recruitment during early embryogenesis. Nat. Cell Biol. 11, 172–182 (2009).

58. Kruszka, P. et al. A CCR4-NOT Transcription Complex, Subunit 1, CNOT1, Variant Associated with Holoprosencephaly. Am. J. Hum. Genet. 104, 990–993 (2019).

59. Zheng, X. et al. Cnot1, Cnot2, and Cnot3 maintain mouse and human ESC identity and inhibit extraembryonic differentiation. Stem Cells 30, 910–922 (2012).

60. Vaites, L. P., Paulo, J. A., Huttlin, E. L. & Harper, J. W. Systematic analysis of human cells lacking ATG8 proteins uncovers roles for gabaraps and the CCZ1/MON1 regulator c18orf8/rmc1 in macroautophagic and selective autophagic flux. Mol. Cell. Biol. 38, (2018).

61. Tsui, C. K. et al. CRISPR-Cas9 screens identify regulators of antibody-drug conjugate toxicity. Nat. Chem. Biol. 15, 949–958 (2019).

62. Zhang, Z. & Gerstein, M. Identification and characterization of over 100 mitochondrial ribosomal protein pseudogenes in the human genome. Genomics 81, 468–480 (2003).

63. Lenoir, G., Williamson, P. & Holthuis, J. C. M. On the origin of lipid asymmetry: the flip side of ion transport. Curr. Opin. Chem. Biol. 11, 654–661 (2007).

64. Uchida, Y. et al. Intracellular phosphatidylserine is essential for retrograde membrane traffic through endosomes. Proc. Natl. Acad. Sci. USA 108, 15846–15851 (2011).

65. Cho, K.-J. et al. Inhibition of Acid Sphingomyelinase Depletes Cellular Phosphatidylserine and Mislocalizes K-Ras from the Plasma Membrane. Mol. Cell. Biol. 36, 363–374 (2016).

66. Andersen, J. P. et al. P4-ATPases as Phospholipid Flippases-Structure, Function, and Enigmas. Front. Physiol. 7, 275 (2016).

67. Huang, H. W., Goldberg, E. M. & Zidovetzki, R. Ceramides modulate protein kinase C activity and perturb the structure of Phosphatidylcholine/Phosphatidylserine bilayers. Biophys. J. 77, 1489–1497 (1999).

68. Kobayashi, T. & Menon, A. K. Transbilayer lipid asymmetry. Curr. Biol. 28, R386–R391 (2018).

69. Takatsu, H. et al. Phospholipid flippase ATP11C is endocytosed and downregulated following Ca2+-mediated protein kinase C activation. Nat. Commun. 8, 1423 (2017).

70. Xie, Z. & Cai, T. Na+-K+--ATPase-mediated signal transduction: from protein interaction to cellular function. Mol Interv 3, 157–168 (2003).

71. Van Der Woerd, W. L., Wichers, C. G. K. & Vestergaard, A. L. Defective plasma membrane targeting of p. I661T-ATP8B1, associated with familial intrahepatic cholestasis, can be rescued in vitro by CFTR correctors. ATP8B1 … (2016).

72. Kato, U. et al. Role for phospholipid flippase complex of ATP8A1 and CDC50A proteins in cell migration. J. Biol. Chem. 288, 4922–4934 (2013).

73. Rosa-Ferreira, C., Christis, C., Torres, I. L. & Munro, S. The small G protein Arl5 contributes to endosome-to-Golgi traffic by aiding the recruitment of the GARP complex to the Golgi. Biol. Open 4, 474–481 (2015).

74. Whyte, J. R. C. & Munro, S. Vesicle tethering complexes in membrane traffic. J. Cell Sci. 115, 2627–2637 (2002).

75. Yu, I.-M. & Hughson, F. M. Tethering factors as organizers of intracellular vesicular traffic. Annu. Rev. Cell Dev. Biol. 26, 137–156 (2010).

76. D’Souza-Schorey, C. & Chavrier, P. ARF proteins: roles in membrane traffic and beyond. Nat. Rev. Mol. Cell Biol. 7, 347–358 (2006).

77. Houghton, F. J. et al. Arl5b is a Golgi-localised small G protein involved in the regulation of retrograde transport. Exp. Cell Res. 318, 464–477 (2012).

78. Pasqualato, S., Renault, L. & Cherfils, J. Arf, Arl, Arp and Sar proteins: a family of GTP-binding proteins with a structural device for “front-back” communication. EMBO Rep. 3, 1035–1041 (2002).

79. Brinkman, E. K., Chen, T., Amendola, M. & van Steensel, B. Easy quantitative assessment of genome editing by sequence trace decomposition. Nucleic Acids Res. 42, e168 (2014).

80. Tepper, A. D. et al. Sphingomyelin hydrolysis to ceramide during the execution phase of apoptosis results from phospholipid scrambling and alters cell-surface morphology. J. Cell Biol. 150, 155–164 (2000).

81. Zhai, X. et al. Phosphatidylserine Stimulates Ceramide 1-Phosphate (C1P) Intermembrane Transfer by C1P Transfer Proteins. J. Biol. Chem. 292, 2531–2541 (2017).

82. Holopainen, J. M., Brockman, H. L., Brown, R. E. & Kinnunen, P. K. Interfacial interactions of ceramide with dimyristoylphosphatidylcholine: impact of the N-acyl chain. Biophys. J. 80, 765–775 (2001).

83. Carrer, D. C. & Maggio, B. Phase behavior and molecular interactions in mixtures of ceramide with dipalmitoylphosphatidylcholine. J. Lipid Res. 40, 1978–1989 (1999).

84. Wu, B. X. et al. Identification of novel anionic phospholipid binding domains in neutral sphingomyelinase 2 with selective binding preference. J. Biol. Chem. 286, 22362–22371 (2011).

85. Hassouneh, L. K. M. Effects of vitamin E on the synthesis of phospholipids and brain functions in old rats. Neurophysiology 50, 166–172 (2018).

